## Supplemental Figures S1-6, Tables S1-S3 for "The canonical ER stress IRE1α/XBP1 pathway mediates skeletal muscle wasting during pancreatic cancer cachexia"

### Targeting the canonical ER stress IRE1α/XBP1 pathway counteracts pancreatic cancer-induced skeletal muscle wasting

by

This file contains Supplemental Figures S1-S6 and Tables S1-S3

### SUPPLEMENTAL FIGURES

#### FIGURE S1

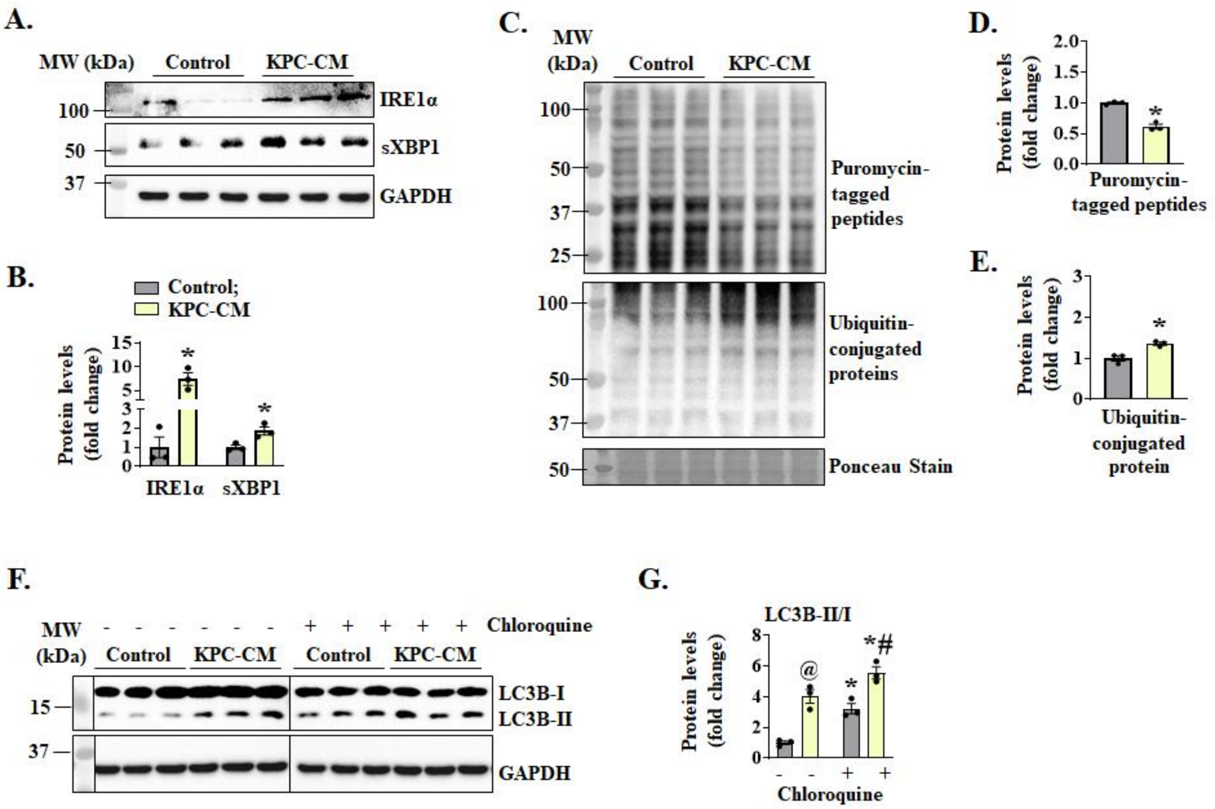


**Figure S1. KPC tumor-derived factors activate IRE1α/XBP1 signaling and regulate protein turnover in cultured myotubes.** Mouse primary myotubes were treated with conditioned media from KPC cells (KPC-CM) or with exhausted differentiation media (as control). After incubation for 24 h, the cells were collected for western blot analysis. **(A)** Immunoblots and **(B)** densitometry analysis of levels of IRE1α and sXBP1 protein in control and KPC-CM treated myotube cultures. **(C)** Immunoblots, and densitometry analysis of levels of **(D)** puromycin-tagged peptides and **(E)** Ubiquitin-conjugated proteins in control and KPC-CM treated cultures. n=3 biological replicates per group. All data are presented as mean ± SEM. *$p < 0.05$, values significantly different from corresponding control cultures; analyzed by unpaired Student *t* test. After 24 h treatment with DM or KPC-CM, 100µM chloroquine or vehicle alone was added to myotube culture media for 1 h, followed by collection of the cells for western blotting. **(F)** Immunoblot and **(G)** quantification of ratio of LC3B-II/I proteins in control and KPC-CM treated myotube cultures treated with vehicle alone or chloroquine. n=3 biological replicates per group. All data are presented as mean ± SEM. *$p < 0.05$, values significantly different from corresponding cultures treated with vehicle alone; @$p<0.05$, values significant different from control cultures treated with vehicle alone. #$p < 0.05$, values significantly different from control cultures treated with chloroquine, analyzed by two-way ANOVA followed by Tukey's multiple comparison test.

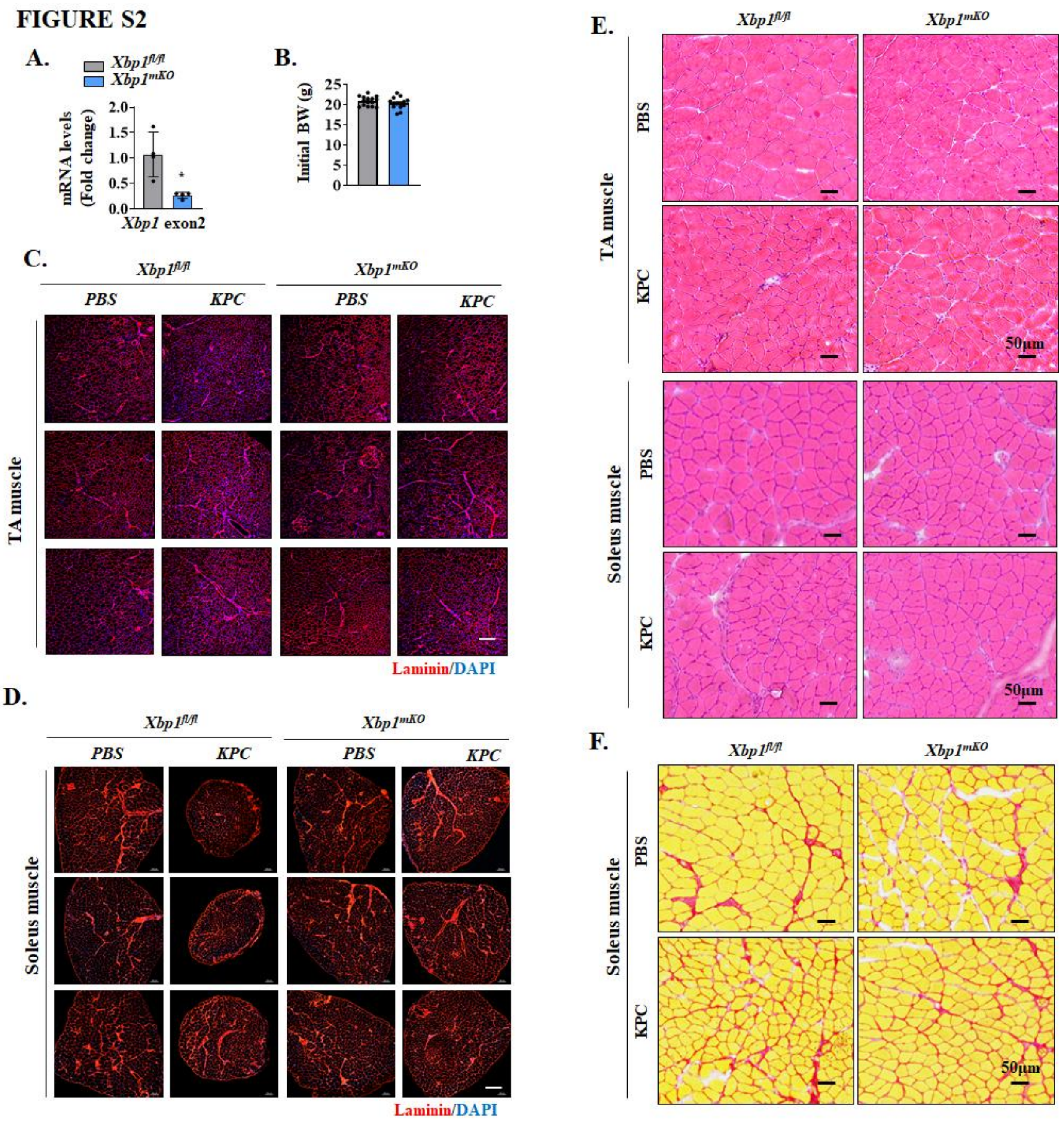


**Figure S2. Targeted deletion of XBP1 inhibits muscle atrophy in KPC tumor-bearing mice.** **(A)** Relative mRNA levels of XBP1 determined by qRT-PCR analysis using primer set specific for Xbp1 exon2, a sequence flanked by the loxP sites, in gastrocnemius (GA) muscle of *Xbp1*$^{fl/fl}$ and *Xbp1*$^{mKO}$ mice. n=4 mice per group. Data are presented as mean ± SEM. *$p$ < 0.05, values significantly different from GA muscle of *Xbp1*$^{fl/fl}$ mice, analyzed by unpaired Student $t$ test. **(B)** Quantification of initial body weight of *Xbp1*$^{fl/fl}$ and *Xbp1*$^{mKO}$ mice. n=14-15 mice per group. No significant difference was observed using unpaired Student *t* test. Transverse sections of TA and soleus muscle isolated from PBS- or KPC cells-injected *Xbp1*$^{fl/fl}$ and *Xbp1*$^{mKO}$ mice were used for anti-laminin and DAPI staining or H&E staining. Anti-laminin and DAPI stained sections of **(C)** TA and **(D)** soleus muscle from multiple mice. Scale bar, 200μm. **(E)** Representative photomicrographs of H&E-stained transverse sections of TA (upper panel) and soleus (lower panel) muscle of control and KPC tumor-bearing *Xbp1*$^{fl/fl}$ and *Xbp1*$^{mKO}$ mice. Scale bar, 50μm. **(F)** Representative photomicrographs of Sirius red-stained soleus muscle sections of *Xbp1*$^{fl/fl}$ and *Xbp1*$^{mKO}$ mice injected with PBS or KPC cells. Scale bar, 50μm.

#### FIGURE S3

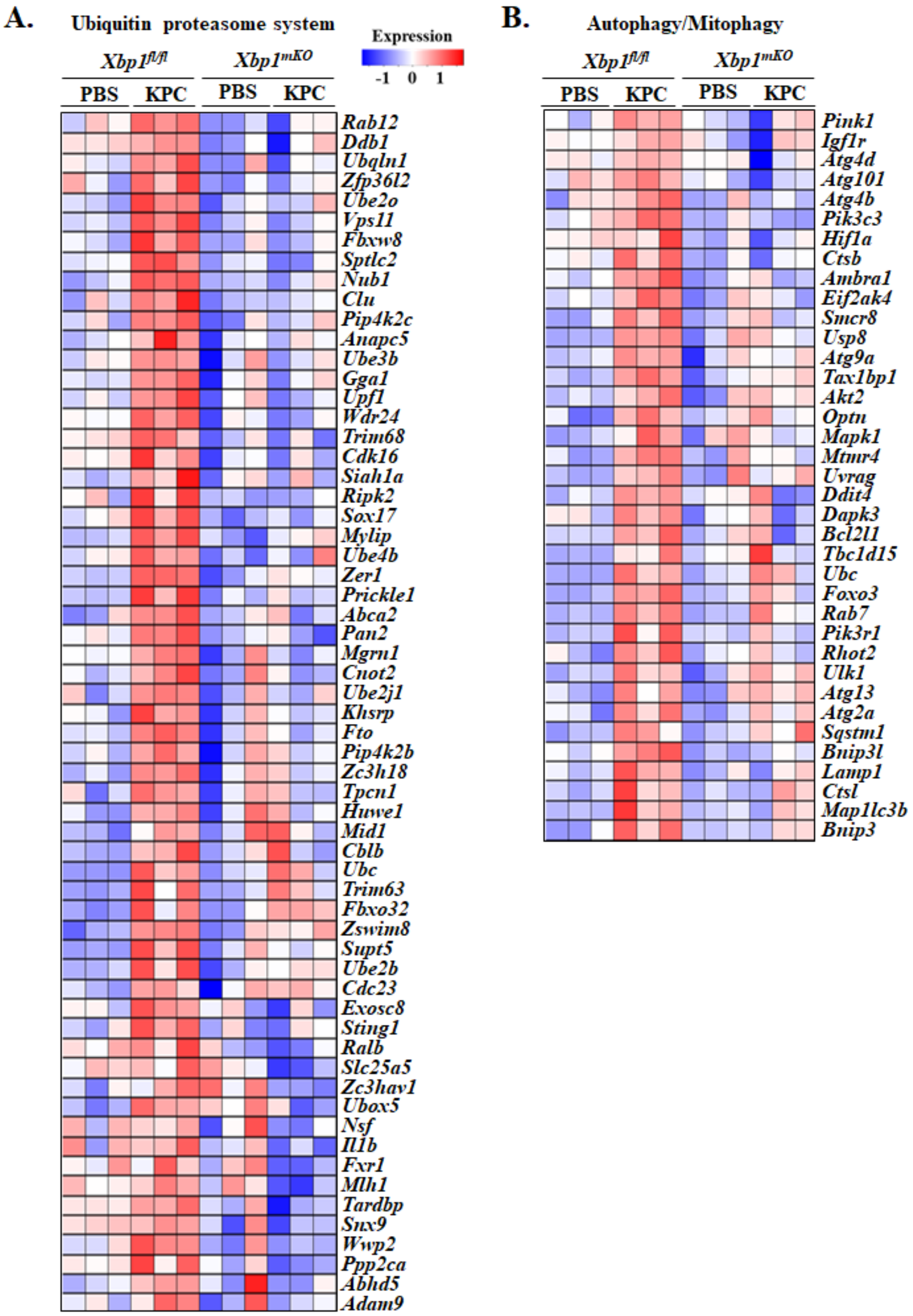


**Figure S3. Targeted ablation of XBP1 inhibits the expression of multiple genes involved in proteolysis.** Heatmap representation of RNA-Seq dataset analysis showing relative expression of genes involved in **(A)** Ubiquitin proteasome system (UPS), and **(B)** autophagy/mitophagy in gastrocnemius muscle of control and KPC tumor-bearing *Xbp1*$^{fl/fl}$ and *Xbp1*$^{mKO}$ mice.

**FIGURE S4**

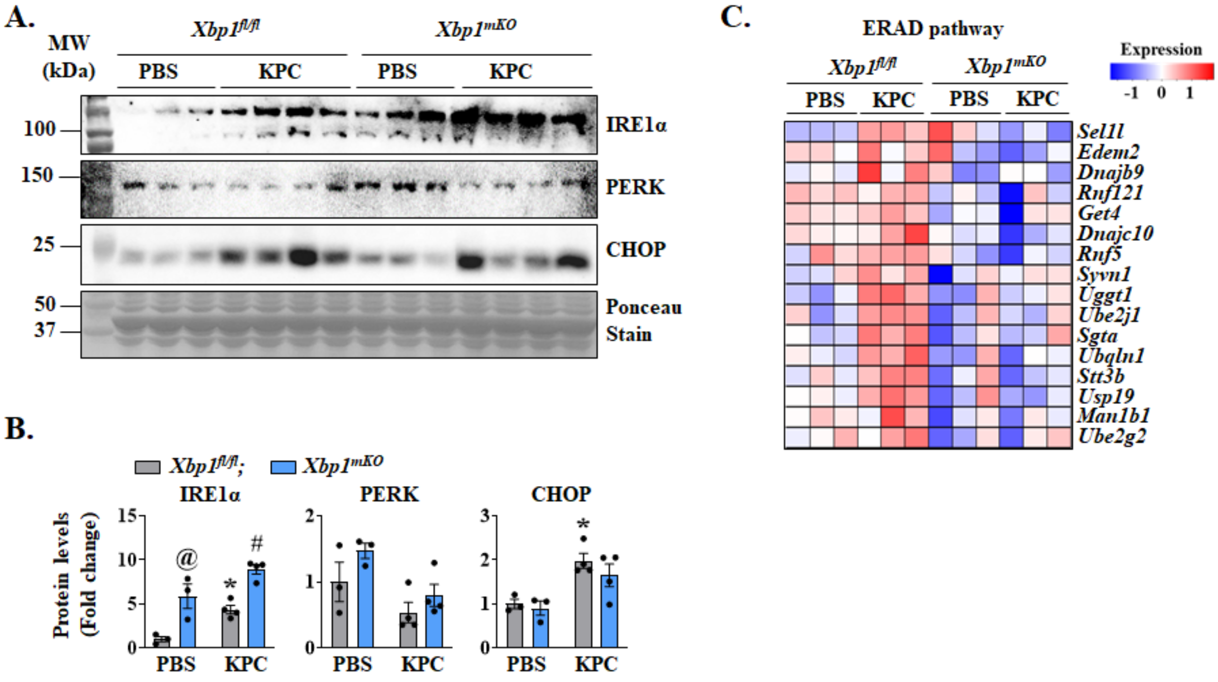


**Figure S4. Effect of targeted ablation of XBP1 on levels of UPR markers. (A)** Immunoblots and **(B)** densitometry analysis of protein levels of IRE1α, PERK, and CHOP in gastrocnemius (GA) muscle of *Xbp1*$^{fl/fl}$ and *Xbp1*$^{mKO}$ mice injected with PBS or KPC cells. n=3-4 mice per group. Data are presented as mean ± SEM. $^{@}p < 0.05$, values significantly different from PBS-injected *Xbp1*$^{fl/fl}$ mice; $^{*}p < 0.05$, values significantly different from corresponding PBS-injected mice; $^{\#}p < 0.05$, values significantly different from KPC tumor-bearing *Xbp1*$^{fl/fl}$ mice, analyzed by two-way ANOVA followed by Tukey's multiple comparison test. **(C)** Heatmap representing relative gene expression of various molecules involved in ER associated degradation (ERAD) pathway in GA muscle of control and KPC tumor-bearing *Xbp1*$^{fl/fl}$ and *Xbp1*$^{mKO}$ mice analyzed using bulk RNA-seq dataset.

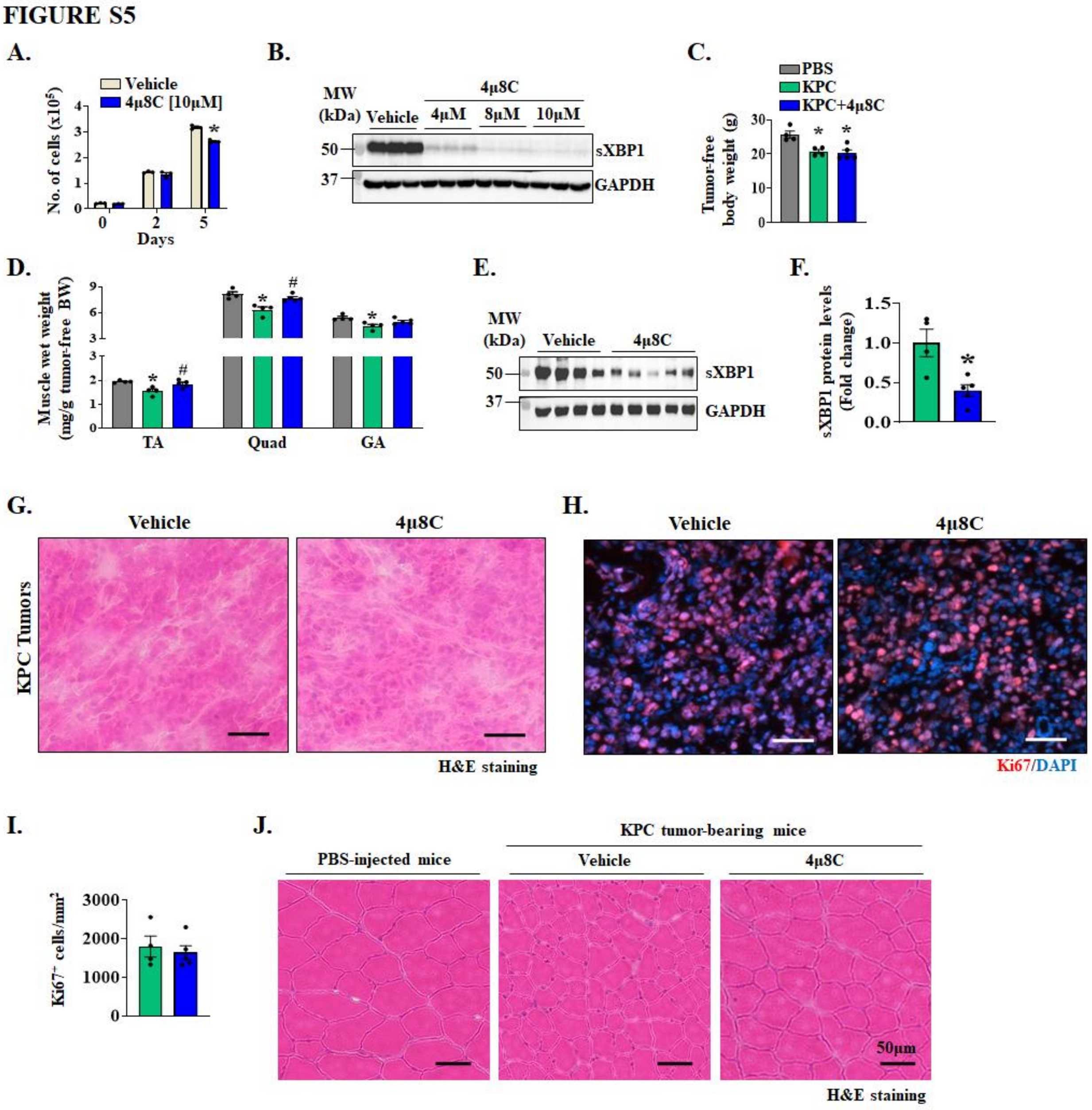


**Figure S5. Effect of pharmacological inhibition of IRE1α/XBP1 axis on cancer cachexia.** **(A)** Quantification of number of proliferating KPC cells on day 0, 2 and 5 after treatment with vehicle alone or 10 µM 4µ8C. n=3 biological replicates per group. Data are presented as mean ± SEM. *$p < 0.05$, values significantly different from vehicle-treated KPC cultures on day 5, analyzed by unpaired Student *t* test. **(B)** Immunoblot showing levels of sXBP1 protein in KPC cells treated with vehicle alone or indicated concentrations of 4µ8C for 24 h. **(C)** Quantification of tumor-free body weight (BW) of control and KPC tumor-bearing mice treated with vehicle alone or 4μ8C after 18 days of KPC cells injection into the pancreas. **(D)** Quantification of wet weight of TA, Quad, and GA muscle normalized by tumor-free BW. n=4-5 mice per group. Data are presented as mean ± SEM. $^{*}p < 0.05$, values significantly different from PBS-injected mice; $^{\#}p < 0.05$, values significantly different from vehicle-treated KPC tumor-bearing mice, analyzed by one-way ANOVA, followed by Tukey's multiple comparison test. **(E)** Immunoblot and **(F)** densitometry analysis showing levels of sXBP1 protein in KPC tumors of mice treated with vehicle alone or 4μ8C. Data are presented as mean ± SEM. $^{*}p < 0.05$, values significantly different from vehicle-treated KPC tumor-bearing mice, analyzed by unpaired Student *t* test. Representative photomicrographs of KPC tumors after **(G)** H&E staining, or **(H)** anti-Ki67 and DAPI staining. Scale bar, 50μm. **(I)** Quantification of number of $Ki67^{+}$ cells per unit area ($mm^2$) in KPC tumors of mice treated with vehicle alone of 4μ8C. No significant difference was observed using unpaired Student *t* test. **(J)** Transverse sections of TA muscle of control and vehicle or 4μ8C-treated KPC tumor-bearing mice were generated and used for H&E staining. Representative photomicrographs of H&E-stained sections are presented here. Scale bar, 50μm.

#### FIGURE S6

##### Fig.1I

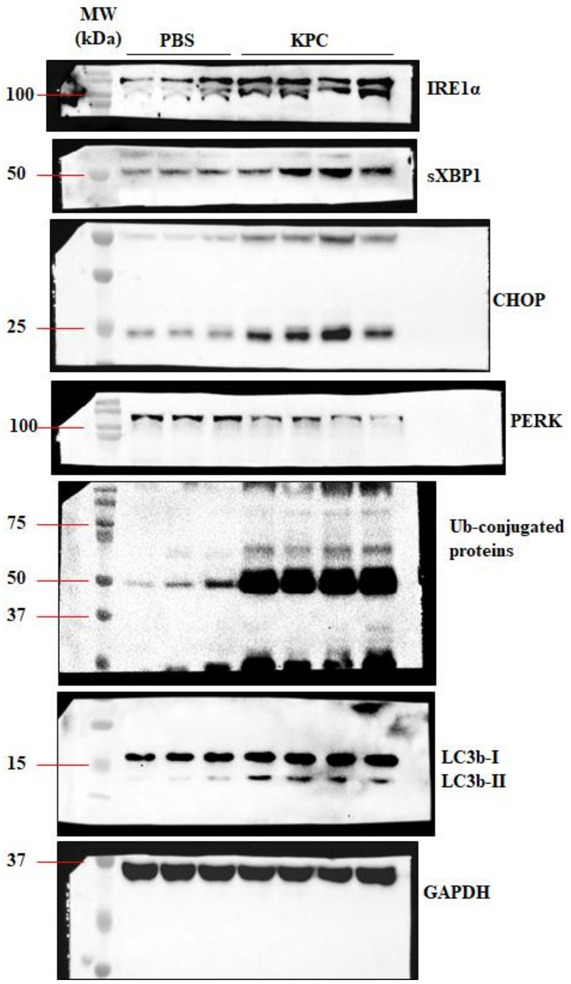


##### Fig.3D

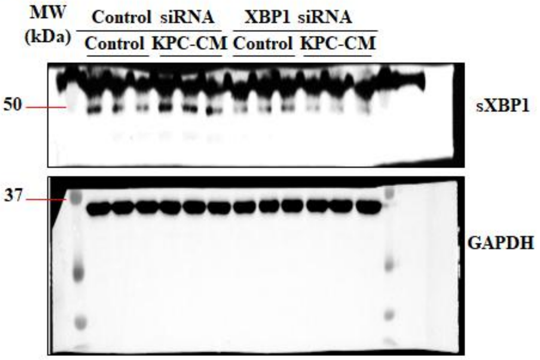


##### Fig.3J

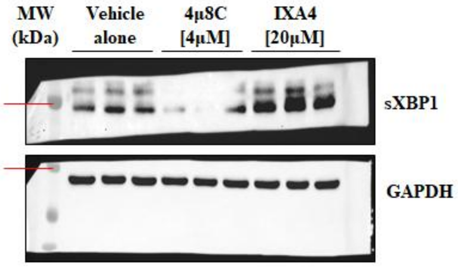

#### FIGURE S6 (continued)

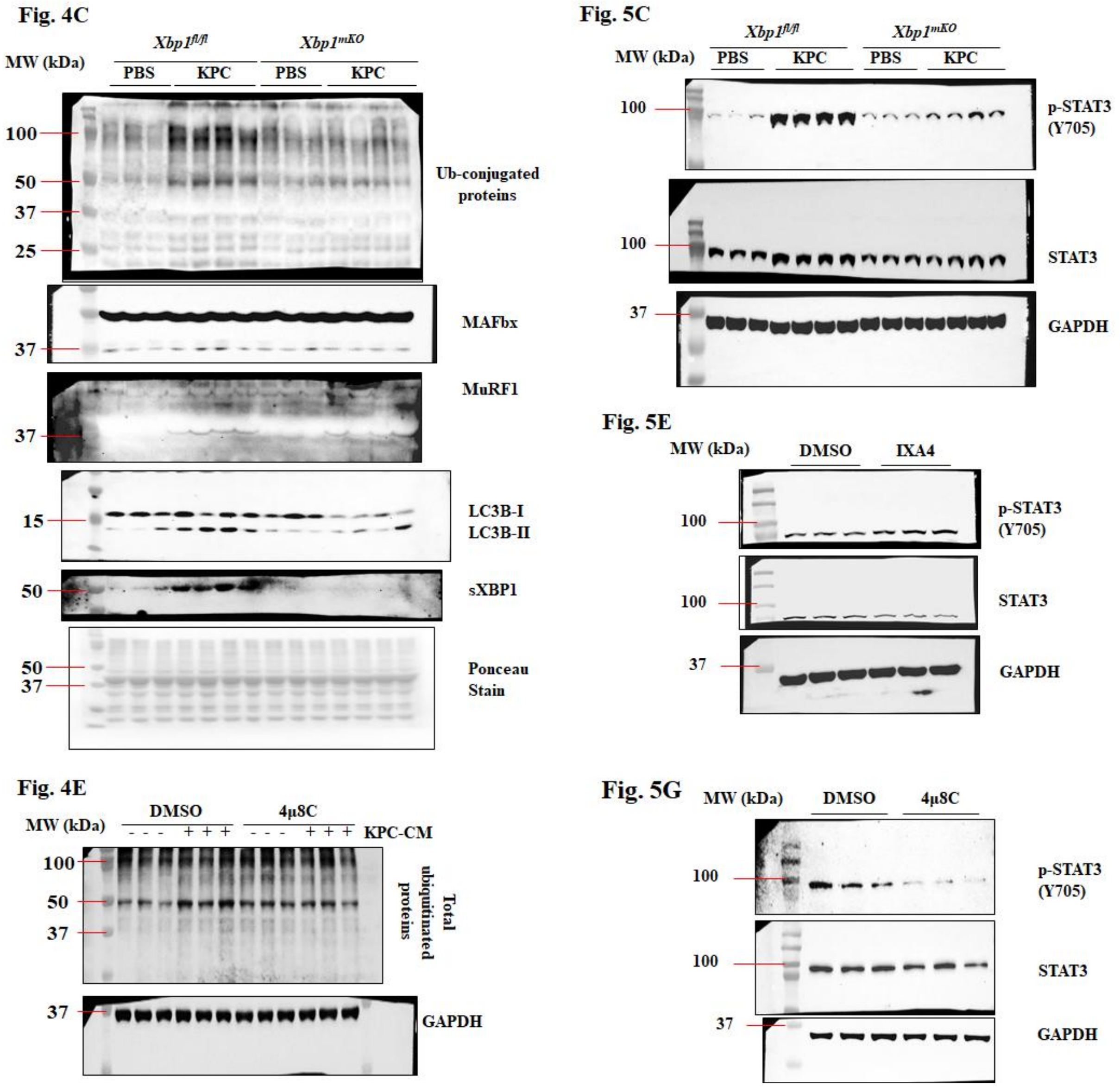

#### FIGURE S6 (continued)

##### Fig. 5K

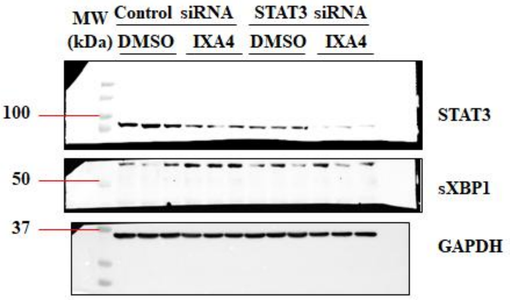


##### Fig. 6F

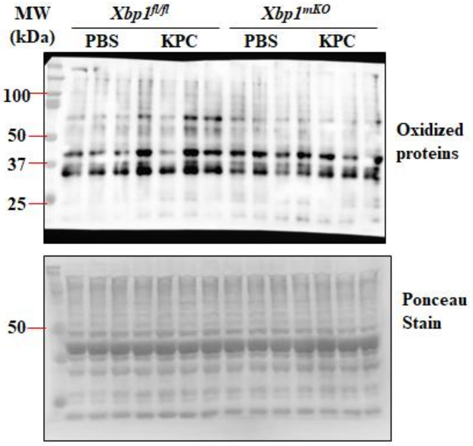


##### Fig. 7C

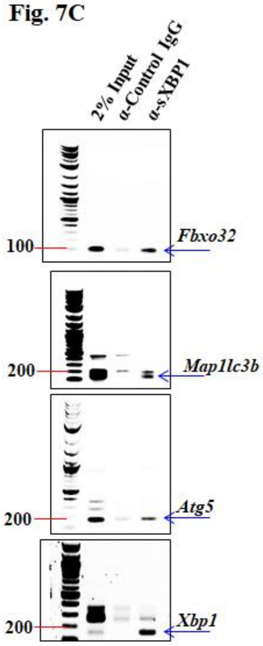


##### Fig. 7G

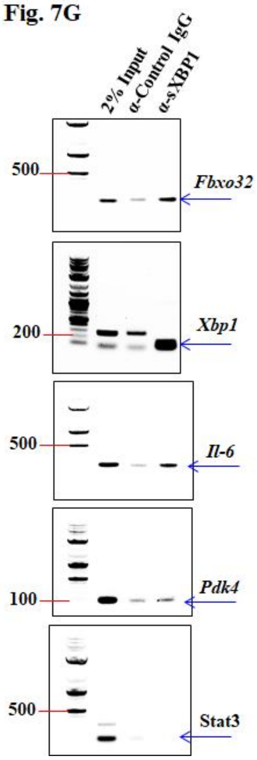


##### Fig. 7E

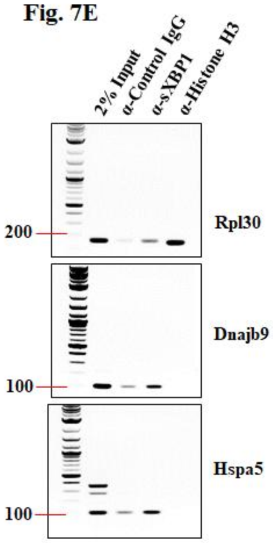


##### Fig. 7I

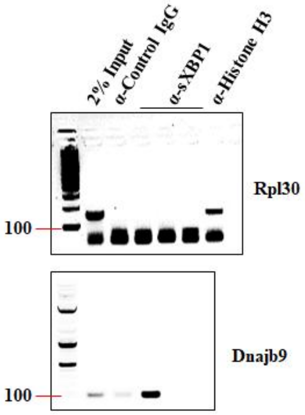

#### FIGURE S6 (continued)

**Fig. 8I**

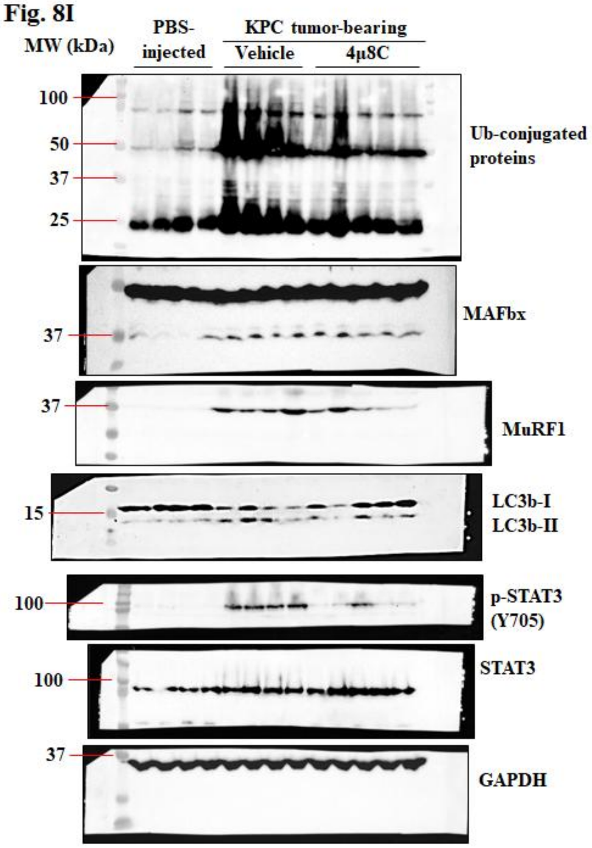


**Fig. 8K**

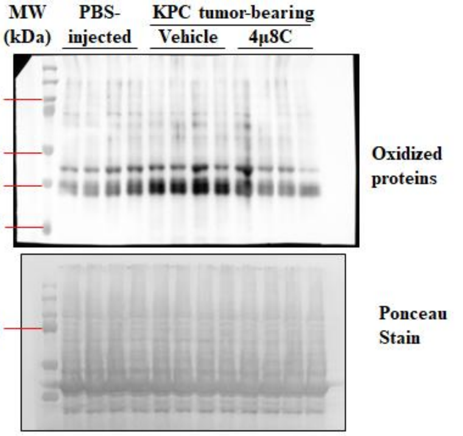


**Fig. S1A**

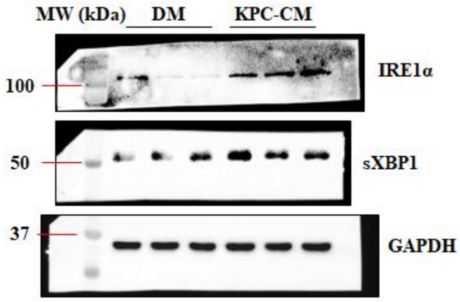


**Fig. S1C**

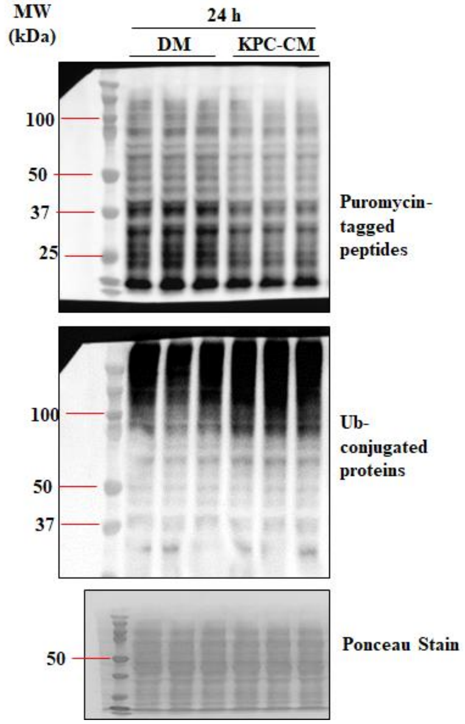

#### FIGURE S6 (continued)

**Fig. S1F**

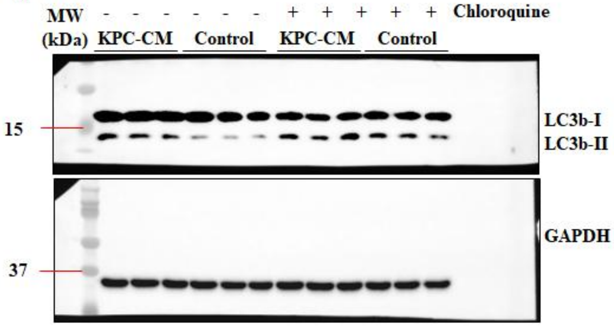


**Fig. S5B**

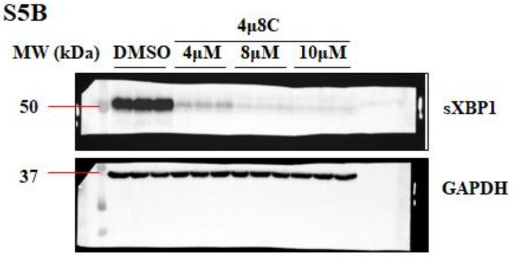


**Fig. S4A**

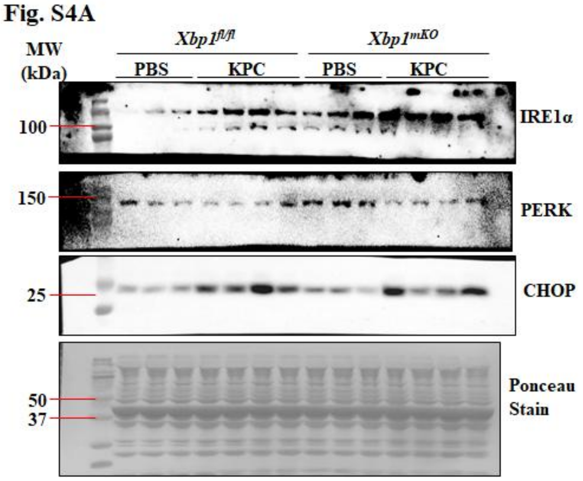


**Fig. S5E**

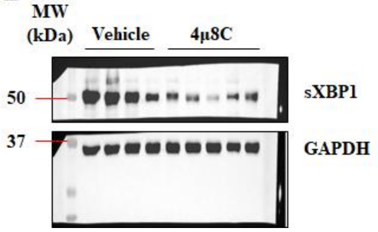


**Figure S6. Uncropped western blot and agarose gel images.**

**Table S1.** List of antibodies used for western blot and immunostaining

| Antibody | Source and Catalog no. | Dilution | Analysis |
|---|---|---|---|
| MAFbx | ECM Biosciences # AP2041 | 1:1000 | WB |
| MuRF1 | R&D Systems #AF5366 | 1:1000 | WB |
| Beclin1 | Cell Signaling # 3495S | 1:1000 | WB |
| LC3b | Cell Signaling #2775S | 1:1000 | WB |
| Ubiquitin | Santa Cruz # sc-8017 | 1:1000 | WB |
| IRE1alpha | Cell Signaling # 3294S | 1:1000 | WB |
| sXBP1 (E9V3E) | Cell Signaling # 40435 | 1:1000 / 1:50 | WB / ChIP |
| PERK | Cell Signaling # 3192 | 1:1000 | WB |
| CHOP | Cell Signaling # 2895S | 1:1000 | WB |
| Phospho-STAT3 (Y705) | Cell Signaling # 9145 | 1:1000 | WB |
| STAT3 | Cell Signaling # 30835 | 1:1000 | WB |
| GAPDH | Cell Signaling # 2118 | 1:1000 | WB |
| Anti-rabbit IgG HRP | Cell Signaling #7074S | 1:2000 | WB |
| Anti-mouse IgG HRP | Cell Signaling #7076S | 1:2000 | WB |
| Anti-goat IgG HRP | Invitrogen #A15999 | 1:2000 | WB |
| Laminin | Sigma Chemical Co. # L9393 | 1:1000 | IF |
| Myosin heavy chain (MyHC) | DSHB # MF20 | 1:100 | IF |
| Perilipin-2 | Proteintech # 15294-1-AP | 1:300 | IF |
| Anti-Mouse IgG2b AF555 | Invitrogen # A21147 | 1:1000 | IF |
| Anti-rabbit IgG AF555 | Invitrogen #A31572 | 1:1000 | IF |

**Table S2.** List of primer sequences used for qPCR and ChIP assays.

| Gene Name | Forward primer (5'-3') | Reverse primer (5'-3') |
|---|---|---|
| Xbp1 (exon2) | CCTGAGCCCGGAGGAGAA | CTCGAGCAGTCTGCGCTG |
| β-actin | CAGGCATTGCTGACAGGATG | TGCTGATCCACATCTGCTGG |
| Hspa5 (ChIP) | TGGTGGCATGGACCAATCAG | CGCCGACTCGCCTTATATAC |
| Dnajb9 (ChIP) | GAGCCGACCTACACGAAAC | AGGACCAAACGGCAACAA |
| Ern1 | CCTTTGCTGATAGTCTCTGCCCAT | TTACCACCAGTCCATCGCCATT |
| Xbp1 | TGTCCATTCCCAAGCGTGTTCT | TGGAGCAGCAAGTGGATTT |
| sXbp1 | AAGAACACGCTTGGGAATGG | CTGCACCTGCTGCGGAC |
| Becn1 | TGAAATCAATGCTGCCTGGG | CCAGAACAGTATAACGGCAACTCC |
| Map1lc3b | CTGGTGAATGGGCACAGCATG | CGTCCGCTGGTAACATCCCTT |
| Atg12 | ACAAAGAAATGGGCTGTGGAGC | GCAGTAATGCAGGACCAGTTTACC |
| Fbxo32 | GTCGCAGCCAAGAAGAGAAAGA | TGCTATCAGCTCCAACAGCCTT |
| Trim63 | TACTGCATCTCCATGCTGGTG | TGGCGTAGAGGGTGTCAAACTT |
| Fbxo30 | TCGTGGAATGGTAATCTTGC | CCTCCCGTTTCTCTATCACG |
| Ddit3 | TGAAAGCAGAACCTGGTCCA | CACTGTTCATGCTTGGTGCA |
| Dnajb9 | TTAGCCATGAAGTACCACCCTGAC | TTCCGACTATTGGCATCCGA |
| Edem1 | CGGCTATGACAACTACATGG | GTTCAGATTGGACTCTC |
| Eif2ak3 | ACTCCTGTCTTGGTTGGGTCTGAT | CGTGCTCCGCTTATTCCTTTCT |
| Bloc1s1 | GCCTACATGAACCAGAGAAAG | GTTCTCCACCATTCCAATCC |
| Pdgfr | CCAAGTCAGGTCCCATTTAC | GGTCTTTCTTCGGCTTCTC |
| Scara3 | GGGTTTCTATGGCTGGTTAG | TGCAGAGACCAGAGTAGTT |
| Sparc | CAACTGCAATTGGGCTTTC | ACCAGTCTCACTTCCTCTAC |
| Ppard | TCCATCGTCAACAAAGACGGG | ACTTGGGCTCAATGATGTCAC |
| Ppargc1a | TGGAGTGACATAGAGTGTGCTGC | CTCAAATATGTTCGCAGGCTCA |
| Cd36 | GGCCAAGCTATTGCGACAT | CAGATCCGAACACAGCGTAGA |
| Acox1 | GGATGGTAGTCCGGAGAACA | AGTCTGGATCGTTCAGAATCAAG |
| Acox2 | CCTTCCTAGACCTGCTTCCC | TGTCCGTCATAACAGCCAAG |
| Acox3 | CTTCTGAGAAACGGGGACAA | GCTCGGTAGGCACTAAGAGG |
| Sirt1 | GACGATGACAGAACGTCACAC | CGAGGATCGGTGCCAATCA |
| Hif1a | TGAGCTTGCTCATCAGTTGC | CCATCTGTGCCTTCATCTCA |
| Hadhb | GCACTTTCGGGTTTGTTG | GTGTGAGCTGGAGTCTTATC |
| Irisin (Fndc5) | CACAGAATATATCGTCCATG | GTCACCTCATCTTTGTTCTT |
| Atg5 | ATCAGACCACGACGGAGCGG | GGCGACTGCGGAAGGACAGA |
| Il6 | CCTTCTTGGGACTGATGCTGG | GCCTCCGACTTGTGAAGTGGT |
| Pdk4 | AAAGGACAGGATGGAAGGAATCA | TTTTCCTCTGGGTTTGCACAT |
| MAFbx (ChIP) | CCTCGGAAAACAAGGCGAG | GTCTCTTTGTTGCCGGAAGA |
| LC3b (ChIP) | GTTAACAGATGCTCGCCCAG | TGTGTGTCTCAGTCCGCAG |

| | | |
|---|---|---|
| Atg5 (ChIP) | TTCCGAGTTCAGGCGCTC | GAACCAGAGTGAACCGCAG |
| Xbp1 (ChIP) | CCCGGGACTACAGGACCA | CCACCACCACCATAGCCA |
| Il6 (ChIP) | CTCATGCTTCTTAGGGCTAGC | GAGTGGGTGGGGCTGATT |
| Pdk4 (ChIP) | AAACAAGGACAAGTCTGGGC | TCACTAGAAAGGCCTGGCAC |

**Table S3.** Average TPM values for PBS-injected *Xbp1*$^{fl/fl}$ mice

| Gene | Avg. TPM | Gene | Avg. TPM | Gene | Avg. TPM | Gene | Avg. TPM | Gene | Avg. TPM |
|---|---|---|---|---|---|---|---|---|---|
| Ubqln4 | 40.560 | Cdk16 | 60.291 | Snx9 | 13.452 | Ctsl | 53.734 | Acox1 | 30.195 |
| Sec23a | 21.400 | Siah1a | 5.633 | Wwp2 | 4.556 | Map1lc3b | 79.470 | Plin5 | 8.484 |
| Ube2j1 | 13.683 | Ripk2 | 1.086 | Ppp2ca | 21.888 | Bnip3 | 150.388 | Cyp4f13 | 6.202 |
| Uggt1 | 4.801 | Sox17 | 6.869 | Abhd5 | 14.324 | Edem2 | 3.000 | Por | 10.225 |
| Hsp90ab1 | 511.050 | Mylip | 6.343 | Adam9 | 18.235 | Dnajb9 | 18.080 | Abcd1 | 8.015 |
| Eif2ak1 | 13.608 | Zer1 | 9.286 | Pink1 | 316.814 | Rnf121 | 10.078 | Acadsb | 46.350 |
| Sel1l | 9.256 | Prickle1 | 1.311 | Igf1r | 1.503 | Get4 | 23.249 | Ppard | 9.168 |
| Sar1a | 63.835 | Abca2 | 3.934 | Atg4d | 33.443 | Dnajc10 | 7.647 | Echdc1 | 8.069 |
| Ube4b | 14.744 | Pan2 | 2.947 | Atg101 | 27.022 | Rnf5 | 17.772 | Acox3 | 6.484 |
| Nploc4 | 19.388 | Mgrn1 | 21.404 | Atg4b | 9.843 | Syvn1 | 7.863 | Adipor2 | 27.850 |
| Dnajb12 | 25.350 | Cnot2 | 11.504 | Pik3c3 | 7.219 | Sgta | 53.460 | Acacb | 42.548 |
| Xbp1 | 28.959 | Khsrp | 6.982 | Hif1a | 4.634 | Usp19 | 49.324 | Crat | 132.295 |
| Dnaja2 | 86.389 | Fto | 31.122 | Ctsb | 92.448 | Man1b1 | 5.761 | Pex7 | 14.007 |
| Sec31b | 5.033 | Pip4k2b | 14.309 | Ambra1 | 5.278 | Ube2g2 | 46.501 | Hacl1 | 6.454 |
| Rad23a | 80.304 | Zc3h18 | 9.096 | Smcr8 | 2.811 | Stat5a | 3.486 | Acad10 | 1.680 |
| Stt3b | 26.017 | Tpcn1 | 15.327 | Usp8 | 5.099 | Stat1 | 5.025 | Gcdh | 16.737 |
| Eif2ak4 | 1.249 | Huwe1 | 14.918 | Atg9a | 45.830 | Mcl1 | 62.713 | Ivd | 65.615 |
| Map2k7 | 27.178 | Mid1 | 3.076 | Tax1bp1 | 30.003 | Csf2rb2 | 0.532 | Slc25a17 | 6.535 |
| Ubxn4 | 21.071 | Cblb | 2.069 | Akt2 | 91.849 | Csf2rb | 0.465 | Klhl25 | 1.580 |
| Rad23b | 40.924 | Ubc | 1111.328 | Optn | 29.451 | Jak3 | 1.893 | Dgat1 | 9.182 |
| Ubqln1 | 19.482 | Trim63 | 127.276 | Mapk1 | 33.787 | Il4ra | 2.008 | Pex5 | 10.411 |
| Rab12 | 152.817 | Fbxo32 | 27.393 | Mtmr4 | 1.942 | Tyk2 | 2.101 | Tysnd1 | 6.776 |
| Ddb1 | 59.740 | Zswim8 | 12.448 | Uvrag | 2.447 | Stam | 4.011 | Etfb | 254.387 |
| Zfp36l2 | 5.323 | Supt5 | 22.064 | Ddit4 | 27.123 | Stat2 | 2.084 | Scp2 | 125.223 |
| Ube2o | 6.575 | Ube2b | 372.150 | Dapk3 | 14.855 | Mtor | 5.761 | Adh5 | 46.870 |
| Vps11 | 8.044 | Cdc23 | 6.832 | Bcl2l1 | 8.863 | Osmr | 1.292 | Eci2 | 61.269 |
| Fbxw8 | 6.031 | Exosc8 | 7.630 | Tbc1d15 | 10.628 | Pik3r1 | 3.342 | Echdc2 | 8.136 |
| Sptlc2 | 3.754 | Sting1 | 2.009 | Foxo3 | 3.185 | Il6ra | 0.754 | Akt1 | 23.419 |
| Nub1 | 15.236 | Ralb | 4.303 | Rab7 | 135.455 | Stat6 | 7.432 | Eci1 | 103.961 |
| Clu | 37.495 | Slc25a5 | 36.903 | Pik3r1 | 3.342 | Aox1 | 4.489 | Mlycd | 48.059 |
| Pip4k2c | 4.647 | Zc3hav1 | 1.664 | Rhot2 | 32.676 | Il6st | 23.087 | Decr1 | 38.108 |
| Anapc5 | 259.659 | Ubox5 | 1.903 | Ulk1 | 18.231 | Stat3 | 11.730 | Auh | 38.259 |
| Ube3b | 48.180 | Nsf | 4.456 | Atg13 | 12.485 | C1qtnf9 | 14.000 | Acad12 | 6.596 |
| Gga1 | 20.818 | Il1b | 0.209 | Atg2a | 4.942 | Pdk4 | 138.439 | | |
| Upf1 | 9.204 | Fxr1 | 203.387 | Sqstm1 | 305.650 | Ppargc1a | 4.292 | | |
| Wdr24 | 4.826 | Mlh1 | 4.466 | Bnip3l | 55.155 | Cpt1b | 115.817 | | |
| Trim68 | 3.692 | Tardbp | 66.167 | Lamp1 | 201.108 | Acadm | 189.071 | | |
